# Effect of intranasal administration of neuroEPO in the histological structure of the olfactory mucosa of rats Wistar

**DOI:** 10.1101/2022.11.01.514298

**Authors:** Ketty Suárez Borrás, Gisselle Fernández Peña, Giselle Puldón Seguí, Carlos Luis Pérez Hernández, Yamila Rodríguez Cruz

## Abstract

**Introduction:** Strokes and neurodegenerative diseases are major global health problems. Not only because they cause high mortality and disability, but to the lack of effective therapies. NeuroEPO, a variant of recombinant human erythropoietin (rHu-EPO) with a low sialic acid content, has shown encouraging results as a potential neuroprotective agent when administered intranasally.

**Objective:** To determine the effect of intranasal administration of NeuroEPO on the histological structure of the olfactory mucosa of Wistar rats.

**Materials and Methods:** An experimental, prospective, and longitudinal study was conducted in Wistar rats. Ten healthy animals were randomly distributed into two groups of five each. The control group received a vehicle (0.3 μl/g/day) and the treated group received NeuroEPO (300 μg/kg/day). Both treatments were administered intranasally for 28 days. The histological characteristics of the olfactory mucosa were evaluated. The medians of the study groups were compared using the Mann-Whitney U test.

**Results:** There were no alterations in the histological characteristics of the olfactory epithelium. However, at the level of the lamina propria in the group treated with NeuroEPO, slight hypertrophy, and hyperplasia of the Bowman’s glands was observed.

**Conclusions:** The administration of the nasal formulation of NeuroEPO did not induce histological alterations of the olfactory mucosa of Wistar rats under the experimental conditions of this research.

## INTRODUCTION

With the increase in life expectancy and population aging, the incidence and prevalence of chronic diseases and their consequences have increased. Diseases associated with the Central Nervous System (CNS), including stroke and neurodegenerative diseases have become a major health problem. Not only because of its morbidity and mortality but also because of the lack of effective treatments. (1–3)

In recent decades, intense research has been carried out in the search for new therapies to treat neurological diseases. Different variants of erythropoietin (EPO) have been evaluated for the treatment of neurological diseases reporting satisfactory and encouraging results (4,5).

NeuroEPO, a variant of EPO with a low sialic acid content, is obtained in Cuba, at the Center for Molecular Immunology (CIM) in the production process of recombinant human EPO (rHu-EPO). Preclinical studies have demonstrated the effects of NeuroEPO on CNS protection in models of Alzheimer’s, stroke, and glutamate-induced excitotoxicity (6–9). In clinical studies, intranasal administration has shown to be safe and well tolerated in healthy volunteers and shows favorable effects with clinical improvement in patients with Parkinson’s disease, Alzheimer’s, and spinocerebellar ataxia type 2 (10–14). In addition, NeuroEPO has beneficial effects on glycemia and reproduction in diabetic rats. (15,16)

In previous studies we reported that intranasal administration of NeuroEPO does not affect the structure of the respiratory mucosa or associated lymphoid tissue (17). However, there is insufficient evidence for the effect of NeuroEPO on the olfactory mucosa.

## OBJECTIVE

To determine the effect of intranasal administration of NeuroEPO on the histological structure of the olfactory mucosa in Wistar rats.

## MATERIALS AND METHODS

The research was executed between 2016 and 2017. An experimental, prospective, and longitudinal study was carried out with the participation of the National Center for the Production of Laboratory Animals (CENPALAB) and the Institute of Basic and Preclinical Sciences Victoria de Girón (ICBP-UCMH). Nulliparous Wistar rats, three weeks old and weighing 150 to 175g were used as animal models. They were maintained in the CENPALAB animal farm in suitable environmental conditions of relative humidity of 40-70%, with regulation of the light/dark cycle of 12/12 hours. The temperature ranged from 19 to 25 °C and they were offered access to food and water on demand (18,19).

### Ethical issues

The experiment was carried out after approval by the Institutional Committee for the Care and Use of Laboratory Animals (CICUAL) of CENPALAB. All procedures with animals were performed by highly qualified personnel and followed the indications of institutional and international guidelines for the use and care of animals in the laboratory. (18,19)

### NeuroEPO formulation

The nasal formulation of NeuroEPO whose Active Pharmaceutical Ingredient (API) is rHu-EPO with low sialic acid content (with PCT / cu2006 / 000001 patents and 20050138 to CIDEM, Havana, Cuba) (20) was developed by the Research Center for the Development of Medicines (CIDEM). Both the NeuroEPO and the vehicle were supplied by the CIM. The vehicle contains all substances (excipients) except API.

Animals were randomly distributed in two study groups of 5 rats each.

1. Control group, intranasal administration of vehicle (0.3μl/g/day)
2. Treated group, intranasal administration of NeuroEPO (300 μg/kg/day)

### Intranasal application (IN)

Both groups of rats received intranasal (IN) treatment for 28 days. The animals were immobilized in a supine position. The administration of nasal formulation was applied slowly (approximately two minutes) with an automatic pipette.

### Processing and staining of nasal structures for histological studies

The animals were euthanized on the 28th and end of the experiment. The nasal structures were removed and washed with 0.9% sodium chloride solution and fixed in neutral 4% formalin for 24 hours. Subsequently, they were subjected to a process of bone decalcification with formic acid for a period of 14 days. Nasal tissue samples were obtained for histopathological examination of the third segment of the rats’ nasal cavity (T3) (21). The samples were processed using the paraffin inclusion technique (22). With a Histo-Line Laboratories MR 300 microtome, sections of 5 μm thickness were made to the nasal tissue included, and 5 slides were made per animal. The staining methods used were: the hematoxylin and eosin technique, the histochemical technique of PAS, and the Mallory trichrome technique.

### Histological study of the olfactory mucosa

Histological slides were examined at various magnifications (100x, 400x, and 1000x) under a Motic BA 210 digital light microscope looking for histopathological changes in the olfactory epithelium and lamina propria. Ten randomly selected fields were observed in each slide. A global assessment was carried out in each field, analyzing aspects related to the shape, size, color and location of the structures, also taking into account possible inflammatory or degenerative changes. (23)

### Morphometric study of the olfactory mucosa

The olfactory mucosa was analyzed at the level of the dorsal meatus of the T3 segment. The height of the epithelium was measured from the basement membrane of the epithelium to its apical surface. The thickness of the lamina propria was measured by looking at the connective tissue underlying the epithelium.

Images randomly taken in 10 fields were digitized on each histological sheet. Motic Images Plus 2.0 software was used to photograph each field. A high-resolution digital camera model Moticam coupled to the microscope was used. The photographs were taken with 10x, 40x, and 100x lenses, a binocular 300 (F.N.20) Widefield projection tube, and 100/20.80 light distribution. Measurements were made on the scanned images using the Image Tool version 3 for Windows measurement software.

### Statistical analysis

Data on epithelium height and lamina propria thickness were collected on a Microsoft Excel sheet. The mean and standard deviation were determined as statisticians of central tendency and dispersion respectively. The groups were compared using the Mann-Whitney U test because the data did not present a normal distribution (according to the Kolmogorov-Smirnov test). Statistically significant differences were considered for p<0.05. All statistical analyses were performed with the GraphPad Prism 7 software for Windows.

The primary data from this study are available from Mendeley Data as an open access to information principle. (24)

## RESULTS

### Histological features of the olfactory mucosa

The epithelium of the olfactory mucosa presented a normal histological structure in the rats of both study groups. No inflammatory infiltrate or signs of fibrosis were observed (Figures 1 and 2).

**Figure 1.**
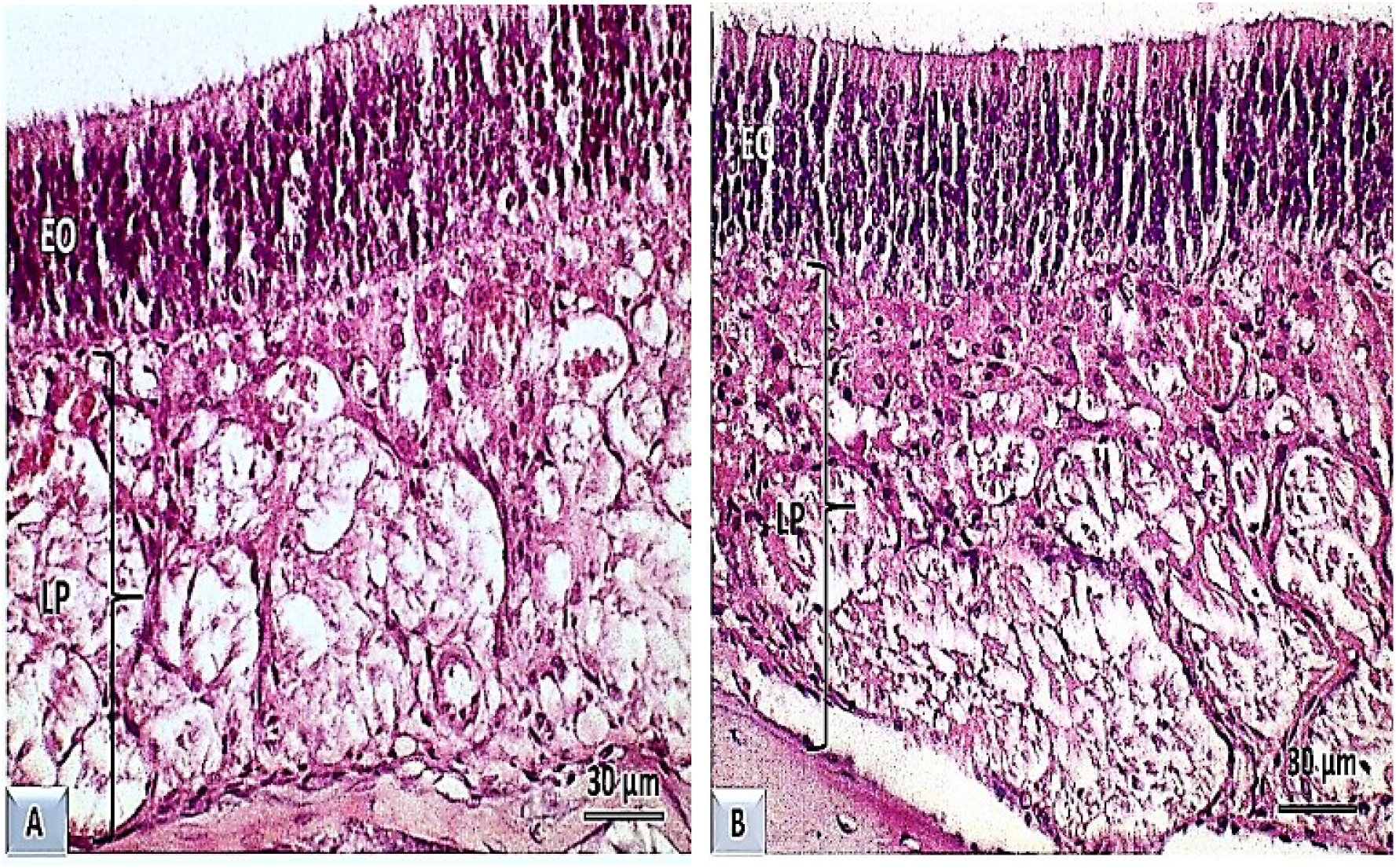
Optical photomicrograph of the olfactory mucosa of Wistar rats. T3. Dorsal meatus. A: Control group. B: Group treated with NeuroEPO. OA, olfactory epithelium. LP: own sheet. Hematoxylin and eosin staining. 400X magnification.

**Figure 2.**
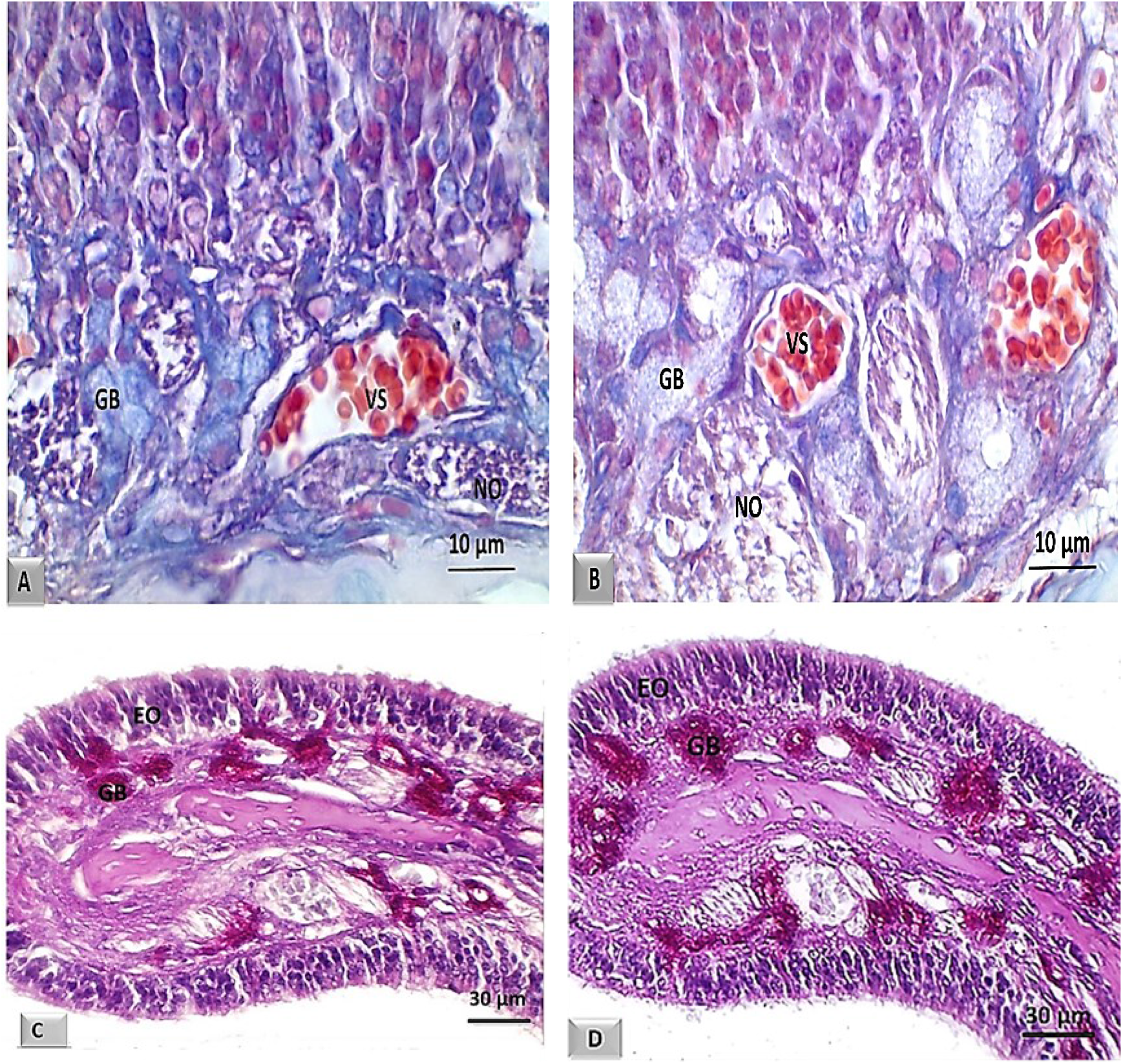
Optical photomicrograph of the olfactory mucosa of Wistar rats. T3. A and C: Control group. B and D: Group treated with NeuroEPO. OA, olfactory epithelium. GB: Bowman’s gland, VS: blood vessels, NO: olfactory nerve. A and B: Mallory trichrome staining. Magnification 1000X. C and D: PAS staining. 400X magnification.

The connective tissue of the lamina propria showed normal characteristics. The olfactory blood vessels and nerves did not present alterations. However, hyperplasia and hypertrophy of Bowman’s glands were observed in all animals in the neuroEPO-treated group (Figures 1B and 2B). The positivity of these glands was evidenced with the histochemical technique of SBP that impresses preserved secretion (Figure 2C and 2D).

### Morphometric study

#### Height of the olfactory epithelium and thickness of the lamina propria

The olfactory mucosa of the rats treated with NeuroEPO showed no differences in relation to the measurements made in the height of the epithelium or the thickness of the lamina propria with respect to that of the rats that received the vehicle (Figure 3).

**Figure 3.**
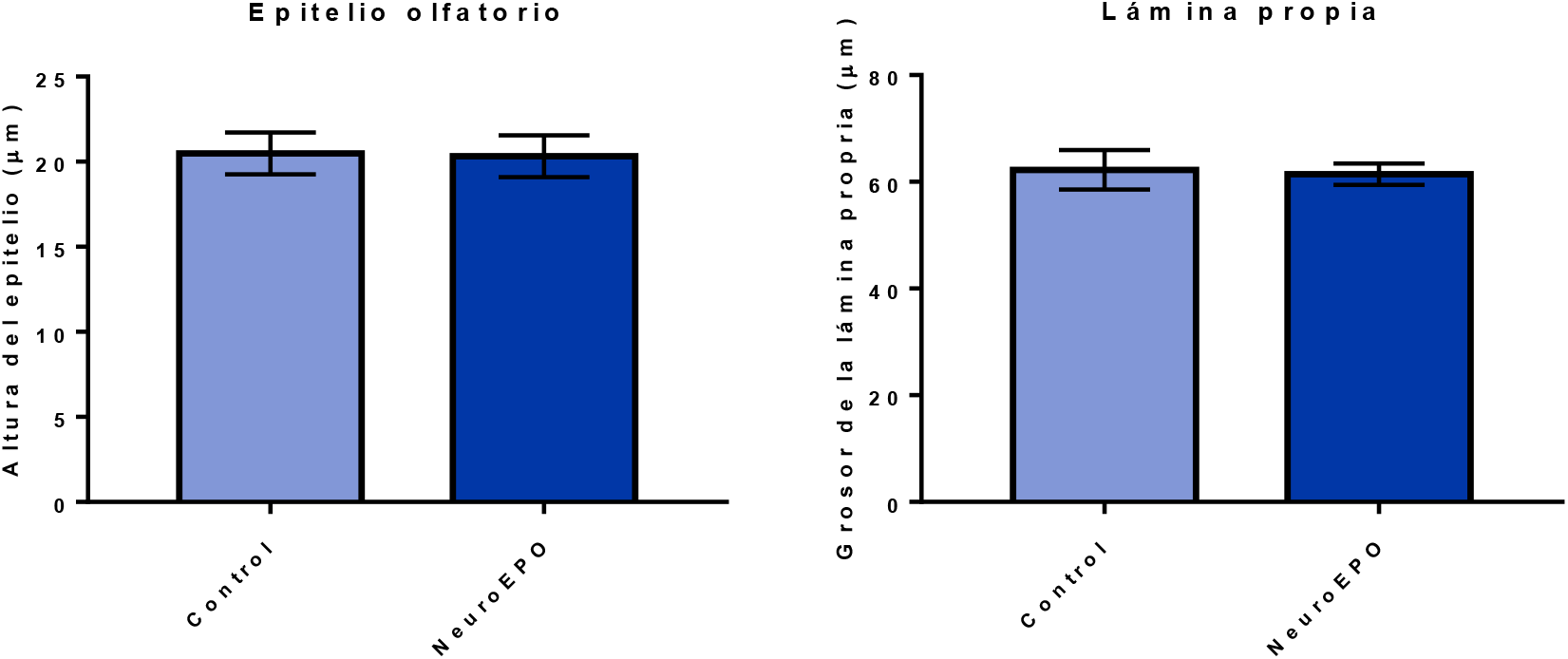
Height of the olfactory epithelium and thickness of the lamina propria of the olfactory mucosa of Wistar rats. Control groups and treated with NeuroEPO. Significance level p >0.05.

## DISCUSSION

Safety is a key issue when designing the formulation of a drug. During the development process, not only the safety of the drug itself but also the active ingredients and excipients within the formulation must be considered (25,26).

The human nasal mucosa has an average physiological pH of 6.3 so it can be considered slightly acidic. Maintaining pH in nasal mucus ensures ciliary clearance function. That is why nasal formulations should have a pH within a range of 4.5 to 6.5 to avoid nasal irritation (25,26).

Not only pH but also osmolarity can induce or favor the presence of toxicological or local effects. However, many substances influence mucociliary clearance through two fundamental mechanisms: by stimulation or by inhibition. Inhibition is the main cause of adverse side effects, such as nasal dryness, irritation, sneezing, nasal itching, rhinitis, and congestion (26).

The present research demonstrated that treatment with intranasal NeuroEPO did not lead to significant histological changes in the olfactory mucosa of Wistar rats, corroborated by the morphometric study. A result that corresponds to a previous study on the respiratory mucosa under the same experimental conditions, in which no histological alterations were evidenced (17).

In any of the cases, leukocyte infiltration, edema, vascular changes, tissue destruction, or connective tissue replacements with vessel proliferation or signs of fibrosis were not observed. Similarly, there were no histological alterations in the cellular or tissue structural characteristics of the olfactory epithelium. These results coincide with research, in which, the use of high doses of intranasal NeuroEPO for 14 days in Wistar rats did not cause signs of inflammatory response or cellular or tissue damage (27).

The results of the current study contrast with several reports. The administration of high doses of substances such as benzalkonium chloride and dantrolene nasally has caused degeneration of the olfactory epithelium, and loss of its cells (both sustaining and receptor cells). In addition, signs of inflammatory response indicative of tissue damage were described (28–31).

In the present study, slight hyperplasia and hypertrophy of the Bowman’s glands was observed in all animals in the treated group. However, it did not affect the thickness of the sheet itself. The positivity of these glands to the histochemical technique of PAS impressed preserved secretion. It is likely that the hypertrophy and hyperplasia of the Bowman’s glands described could be related to reversible adaptive changes of the cells of the glandular epithelium, which does not affect their function and does not translate pathological damage caused by the drug administered nasally.

These results differ from some research in which morphological changes in the lamina propria have been reported with the use of drugs at high doses through the nasal. Absence of bundles of nerve fibers has been demonstrated, as well as reversible and irreversible signs of cell damage in the cells of the glandular epithelium (25,26,31).

In the morphometric study of the olfactory mucosa, no significant differences were observed in the height of the olfactory epithelium and the thickness of the lamina propria in the study groups. In the literature reviewed, no morphometric data of the olfactory mucosa related to the use of NeuroEPO were found. However, a marked decrease in the height of the olfactory epithelium has been reported following the use of some drugs such as methyl bromide and methimazole (25,26,31). In relation to the thickness of the lamina propria, no reports were found in the literature consulted that could be compared with the present study.

This research was limited to the assessment of the structure of the olfactory mucosa at the level (T3) of the nasal cavity so we considered that further studies examining all three levels and including functional assessments could provide stronger evidence.

## CONCLUSIONS

The administration of 300 μg/kg or 6900 IU/kg of NeuroEPO to Wistar rats, for 28 days, did not cause significant structural changes of the olfactory mucosa in our experimental conditions. These results suggest that NeuroEPO can be used intranasally without risk of producing local adverse effects.

Conflict of interest

The authors declare that there is no conflict of interest regarding the results of this publication.

## Notes

### Competing Interest Statement

The authors have declared no competing interest.

## BIBLIOGRAPHY

1. WHO. World health statistics 2022 (Monitoring health of the SDGs) [Internet]. 2022. 1–131 p. Available from: http://apps.who.int/bookorders.

2. Deuschl G, Beghi E, Fazekas F, Varga T, Christoforidi KA, Sipido E, et al. The burden of neurological diseases in Europe: an analysis for the Global Burden of Disease Study 2017. Lancet Public Health. 2020 Oct 1;5(10):e551–67.

3. Feigin VL, Vos T, Nichols E, Owolabi MO, Carroll WM, Dichgans M, et al. The global burden of neurological disorders: translating evidence into policy. Lancet Neurol. 2020 Mar 1;19(3):255–65.

4. Vittori DC, Chamorro ME, Hernández Y v., Maltaneri RE, Nesse AB. Erythropoietin and derivatives: Potential beneficial effects on the brain. J Neurochem [Internet]. 2021 Sep 1 [cited 2022 Oct 24];158(5):1032–57. Available from: https://onlinelibrary.wiley.com/doi/full/10.1111/jnc.15475

5. Rey F, Balsari A, Giallongo T, Ottolenghi S, di Giulio AM, Samaja M, et al. Erythropoietin as a Neuroprotective Molecule: An Overview of Its Therapeutic Potential in Neurodegenerative Diseases. https://doi.org/101177/1759091419871420 [Internet]. 2019 Aug 26 [cited 2022 Oct 24];11. Available from: https://journals.sagepub.com/doi/full/10.1177/1759091419871420

6. Garzón F, Coimbra D, Parcerisas A, Rodriguez Y, García JC, Soriano E, et al. NeuroEPO Preserves Neurons from Glutamate-Induced Excitotoxicity. Journal of Alzheimer’s Disease. 2018 Jan 1;65(4):1469–83.

7. Fernando G, Yamila R, Cesar GJ, Ramón R. Neuroprotective Effects of neuroEPO Using an In Vitro Model of Stroke [Internet]. Behavioral Sciences Multidisciplinary Digital Publishing Institute; Feb 13, 2018 p. 26. Available from: https://www.mdpi.com/2076-328X/8/2/26/htm

8. Maurice T, Mustafa MH, Desrumaux C, Keller E, Naert G, García-Barceló MDLC, et al. Intranasal formulation of erythropoietin (EPO) showed potent protective activity against amyloid toxicity in the Aβ25-35 non-transgenic mouse model of Alzheimer’s disease. https://doi.org/101177/0269881113494939 [Internet]. 2013 Jun 26 [cited 2022 Oct 24];27(11):1044–57. Available from: https://journals.sagepub.com/doi/abs/10.1177/0269881113494939

9. Cruz YR, Strehaiano M, Rodríguez Obaya T, Rodríguez JCG, Maurice T. An Intranasal Formulation of Erythropoietin (Neuro-EPO) Prevents Memory Deficits and Amyloid Toxicity in the APP Swe Transgenic Mouse Model of Alzheimer’s Disease. Journal of Alzheimer’s Disease. 2017 Jan 1;55(1):231–48.

10. Pérez L, Sosa S, Bringas G, López D, Valenzuela C, Peñalver AI, et al. NeuroEPO in mild-to-moderate Alzheimer’s disease. Alzheimer’s & Dementia [Internet]. 2020 Dec [cited 2022 Aug 8];16(S9):e036167. Available from: https://onlinelibrary.wiley.com/doi/full/10.1002/alz.036167

11. Bringas Vega ML, Pedroso Ibáñez I, Razzaq FA, Zhang M, Morales Chacón L, Ren P, et al. The Effect of Neuroepo on Cognition in Parkinson’s Disease Patients Is Mediated by Electroencephalogram Source Activity. Front Neurosci. 2022;0(June):952.

12. Santos-Morales O, Díaz-Machado A, Jiménez-Rodríguez D, Pomares-Iturralde Y, Festary-Casanovas T, González-Delgado CA, et al. Nasal administration of the neuroprotective candidate NeuroEPO to healthy volunteers: A randomized, parallel, open-label safety study. BMC Neurol [Internet]. 2017 Jul 4 [cited 2022 Aug 8];17(1):1–9. Available from: https://link.springer.com/articles/10.1186/s12883-017-0908-0

13. Pedroso I, Garcia M, Casabona E, Morales L, Bringas ML, Pérez L, et al. Protective Activity of Erythropoyetine in the Cognition of Patients with Parkinson’s Disease. Behavioral Sciences 2018, Vol 8, Page 51 [Internet]. 2018 May 21 [cited 2022 Oct 24];8(5):51. Available from: https://www.mdpi.com/2076-328X/8/5/51/htm

14. Rodriguez-Labrada R, Ortega-Sanchez R, Hernández Casaña P, Santos Morales O, Padrón-Estupiñan M del C, Batista-Nuñez M, et al. Erythropoietin in Spinocerebellar Ataxia Type 2: Feasibility and Proof-of-Principle Issues from a Randomized Controlled Study. Movement Disorders [Internet]. 2022 Jul 1 [cited 2022 Oct 24];37(7):1516–25. Available from: https://onlinelibrary.wiley.com/doi/full/10.1002/mds.29045

15. Fernández Romero T, Clapés Hernández S, Pérez Hernández CL, Barreto López JJ, Fernández Peña G, Fernández Romero T, et al. Efecto hipoglicemiante de la NeuroEPO en ratas con y sin diabetes mellitus. Revista Habanera de Ciencias Médicas [Internet]. 2022 [cited 2022 Aug 6];21(1). Available from: http://scielo.sld.cu/scielo.php?script=sci_arttext&pid=S1729-519X2022000100003&lng=es&nrm=iso&tlng=es

16. Fernández Romero T, Clapes Hernández S, Pérez Hernández CL, Núñez López N, Suárez Román G, Fernández Peña G. Protective effect of NeuroEPO on the reproduction of diabetic rats. Revista Habanera de Ciencias Médicas [Internet]. 2022 Sep 22 [cited 2022 Oct 13];21(4):4797. Available from: http://www.revhabanera.sld.cu/index.php/rhab/article/view/4797

17. Suárez Borrás K, Fernández Peña G, Rodríguez Cruz Y, Puldón Seguí G. Intranasal administration of NeuroEPO does not affect the structure of respiratory mucosa in Wistar rats. Revista Habanera de Ciencias Médicas [Internet]. 2022 Sep 22 [cited 2022 Oct 26];21(4):4849. Available from: http://www.revhabanera.sld.cu/index.php/rhab/article/view/4849

18. Couto M, Cates C. Laboratory Guidelines for Animal Care. In Humana, New York, NY; 2019 [cited 2020 Mar 30]. p. 407–30. Available from: http://link.springer.com/10.1007/978-1-4939-9009-2_25

19. McCormick-Ell J, Connell N. Laboratory Safety, Biosecurity, and Responsible Animal Use. ILAR J [Internet]. 2019 Aug 16 [cited 2020 Mar 30];60(1):24–33. Available from: https://academic.oup.com/ilarjournal/advance-article/doi/10.1093/ilar/ilz012/5550511

20. Muñoz Cernada A, García Rodríguez JC, Nuñez Figueredo Y, Pardo Ruiz Z, García Selman JD, Sosa Testé I, et al. Formulaciones nasales de EPORH con bajo contenido de ácido siálico para el tratamiento de enfermedades del sistema nervioso central [Internet]. 2007 [cited 2022 Oct 24]. Available from: https://patentscope.wipo.int/search/es/detail.jsf?docId=WO2007009404

21. Humason GL. Animal tissue techniques. [Internet]. Animal tissue techniques. San Francisco (& London): W. H. Freeman and Company; 1962 [cited 2022 Oct 26]. Available from: https://www.cabdirect.org/cabdirect/abstract/19622204447

22. Uraih LC, Maronpot RR. Normal histology of the nasal cavity and application of special techniques. Environ Health Perspect. 1990 Apr;85:187–208.

23. Kumar V, Abbas AbulK, Aster JonC. Robbins and Cotran Pathologic Basis of Disease [Internet]. 10th ed. Kumar V, K Singh M, editors. Elsevier. Philadelphia: Thomas press India Ltd, Elsevier Publication; 2018 [cited 2022 Oct 26]. Available from: https://www.elsevier.com/books/robbins-and-cotran-pathologic-basis-of-disease/kumar/978-0-323-53113-9

24. Suárez K, Fernández G, Puldón G, Rodriguez Y, Pérez CL. Effect of intranasal administration of neuroEPO in the histological structure of the olfactory mucosa of rats Wistar. [Internet]. Mendeley Data. Mendeley; [cited 2022 Oct 25]. Available from: https://data.mendeley.com/datasets/gc77wttd9h

25. Graff CL, Pollack GM. Nasal Drug Administration: Potential for Targeted Central Nervous System Delivery. J Pharm Sci. 2005 Jun 1;94(6):1187–95.

26. Keller LA, Merkel O, Popp A. Intranasal drug delivery: opportunities and toxicologic challenges during drug development. Drug Deliv Transl Res [Internet]. 2022 Apr 1 [cited 2022 Oct 26];12(4):735–57. Available from: https://link.springer.com/article/10.1007/s13346-020-00891-5

27. Lagarto A, Bueno V, Guerra I, Valdés O, Couret M, López R, et al. Absence of hematological side effects in acute and subacute nasal dosing of erythropoietin with a low content of sialic acid. Experimental and Toxicologic Pathology. 2011 Sep 1;63(6):563–7.

28. Kovalchuk Nataliia. ORGAN-SPECIFIC CONTRIBUTION OF P450 ENZYMES TO BIOACTIVATION AND ACUTE RESPIRATORY TRACT TOXICITY OF NAPHTHALENE by Nataliia Kovalchuk A Dissertation. New York: State University of New York; 2017.

29. Cüreoğlu S, Akkuş M, Osma Ü, Yaldiz M, Oktay F, Can B, et al. The effect of benzalkonium chloride on rabbit nasal mucosa in vivo: an electron microscopy study. European Archives of Oto-Rhino-Laryngology 2002 259:7 [Internet]. 2002 [cited 2022 Oct 26];259(7):362–4. Available from: https://link.springer.com/article/10.1007/s00405-002-0458-x

30. Jiang B, Shi Y, Abou MB, Xu L, Liang G, Wei H. Effects of chronic intranasal dantrolene on nasal mucosa morphology in mice. Eur Rev Med Pharmacol Sci. 2022;26(1):198–203.

31. Xie F, Zhou X, Genter MB, Behr M, Gu J, Ding X. The Tissue-Specific Toxicity of Methimazole in the Mouse Olfactory Mucosa Is Partly Mediated through Target-Tissue Metabolic Activation by CYP2A5. Drug Metabolism and Disposition [Internet]. 2011 Jun 1 [cited 2022 Oct 26];39(6):947–51. Available from: https://dmd.aspetjournals.org/content/39/6/947

